# Plasmid dependent phage eliminate pathogenic bacteria and antibiotic resistance plasmids from the chicken gut microbiome

**DOI:** 10.1101/2025.10.26.684609

**Authors:** Jun Li, Taoran Fu, Xing Ji, Ran Wang, Michael A. Brockhurst, R. Craig MacLean, Tao He

## Abstract

Conjugative plasmids are a key reservoir of antimicrobial resistance (AMR) in commensal and pathogenic bacteria within the gut microbiome. Plasmid-dependent phage (PDPs) are a promising therapeutic option directly targeting AMR plasmids, but the efficacy of PDP therapy *in vivo* is poorly understood. Here, we show that treatment with the PRD1 phage eliminated *Salmonella enterica* carrying a broad-host range IncP AMR plasmid (RP4) from the chicken gut microbiome. Moreover, PRD1 treatment prevented the transfer of the RP4 plasmid to other members of the gut microbiome, whilst causing only minimal disruption to microbiome composition. Our results demonstrate that PDP therapy rapidly eliminates AMR from a natural gut microbiome by killing pathogenic bacteria and blocking AMR plasmid transmission to commensals, highlighting the conjugative pilus as a promising antimicrobial target.

## Introduction

Phage therapy is a promising strategy for treating infections caused by antibiotic resistant bacteria^1,2^. Many of the most clinically important antimicrobial resistance (AMR) genes are carried by conjugative plasmids^3-7^, which readily transfer between pathogenic and commensal bacteria within densely populated diverse microbial communities, exemplified by the gut microbiome^8-12^. Here, the same AMR plasmid can be carried by multiple species, creating a serious challenge for conventional phage therapy, since most lytic phage infect only a limited subset of strains within any given bacterial species^13,14^. An alternative therapeutic approach is to use lytic plasmid-dependent phage (PDPs) that infect bacterial cells using the plasmid conjugative pilus as a receptor^15-21^. Crucially, PDPs can infect the breadth of bacterial species that act as hosts for the targeted plasmid^17,22,23^. As such, PDPs could be a powerful therapeutic option because they combine direct killing of antibiotic resistant pathogens with blocking transmission of AMR plasmids to commensals within complex microbiomes ^15-17^, thus potentially eliminating a key reservoir of AMR.

Poultry production is a major consumer of antibiotics ^24-26^ and acts as an important source of AMR ^27-29^ that can be transmitted to humans through the food chain, contamination of the environment, and the use of chicken manure as a fertilizer ^30^. Consequently, there is an urgent need both to discover alternatives to antibiotics for poultry to reduce antibiotic use, and to develop interventions that reduce the AMR burden arising from poultry farming. Conventional phage therapy can be effective at reducing pathogen abundance in the chicken gut^31^. A previous study showed that the MS2 PDP can target *Salmonella* carrying an F plasmid with a de-repressed conjugative system in the chicken gut^16^. However, AMR plasmids typically show much lower levels of pilus expression than the derepressed F plasmid, and as such it remains unclear whether PDB therapy can be effective against natural AMR plasmids *in vivo*. In addition, it is unknown whether PDPs can effectively block AMR plasmid transmission in a complex gut microbiome and the extent to which PDP therapy perturbs the gut microbiome.

Here, we compared conventional phage therapy against PDP therapy in chickens infected with *Salmonella typhimurium* carrying RP4, a natural broad-host range IncP multidrug resistance plasmid (Figure 1). Chickens were treated with PRD1, a broad-host range lytic phage that infects cells via the IncP conjugative pilus^22,32-35^, or pLT2, a lytic Felix-O1 phage that infects *Salmonella* using LPS as a receptor^36^ or both phages in a cocktail, enabling us to disentangle the effect of targeting the RP4 plasmid versus targeting *Salmonella per se*. As a further control we treated chickens with ampicillin, which selects for the RP4 plasmid, allowing us to measure the impact of phage in comparison to an antibiotic that is widely used in agriculture. We quantified the abundances of *Salmonella*, RP4, and phage over time, and assessed the impact of the phage treatments on chicken gut microbiome composition. Only the treatments containing PRD1 eliminated both *Salmonella* and RP4 from the chicken gut microbiome, whilst also causing no lasting disruption to bacterial community structure.

**Figure 1:**
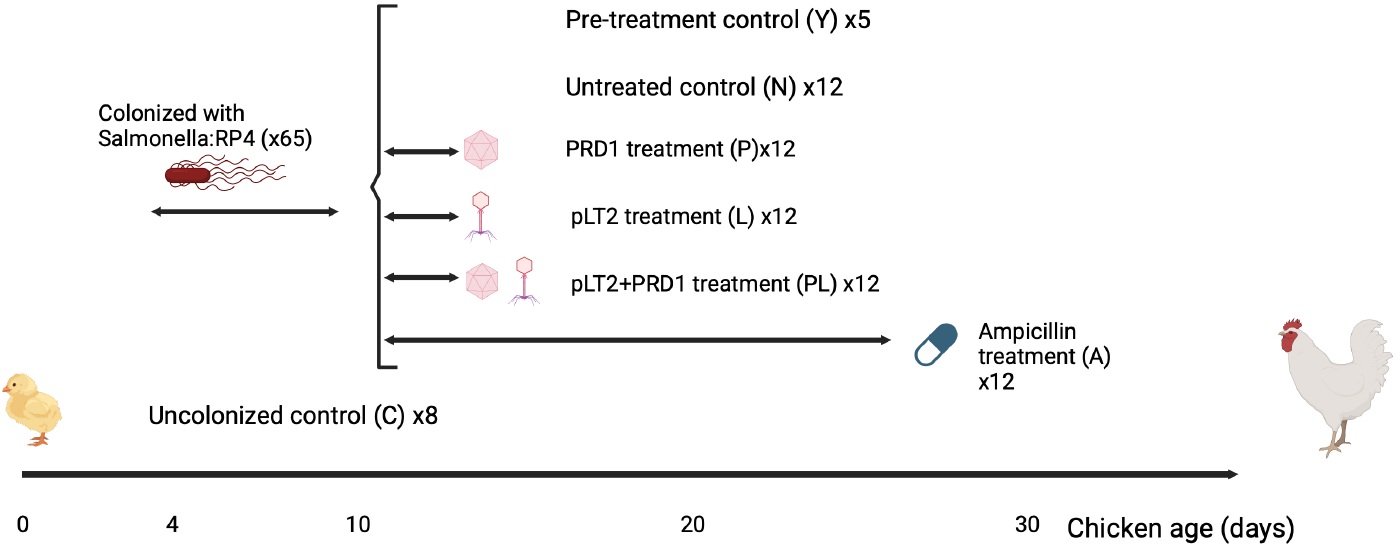
Experimental design. Chicks were colonized with the *Salmonella*:RP4 host strain between day 4 and day 10 or were left as a uncolonized controls. Colonized chicks were treated with PRD1 phage or the pLT2 phage, either alone or in combination, between day 11 and day 13. The responses to phage treatment were compared to untreated control chickens (negative control) and chickens that were treated with ampicillin between day 11 and day 25, which selects for the RP4 plasmid. Chickens were sacrificed at regular time intervals and the density of the host strain, RP4 plasmid, and phage were determined over time. The gut microbiome of a subset of chickens was profiled by shotgun metagenomic sequencing.

Finally, we assembled RP4 plasmid sequences from metagenomic reads to detect mutations selected in response to PDP therapy. Consistent with PRD1 selecting against conjugation *in vivo*, we observed parallel mutations in *traI*, encoding an essential component of RP4’s conjugation machinery, exclusively in PRD1-treated chicken microbiomes.

## Methods and materials

### Plasmid, bacterial strains and phages

The RP4 plasmid contains resistance genes including *bla*TEM-2, *aph(3’)-Ib, tet*(A), and *catA2*. The plasmid was labelled with a *gfp* gene and transformed into the nalidixic acid-resistant *Salmonella Typhimurium* LT2 (see supplementary methods for plasmid construction; Accession PX306537). The PRD1 phage was isolated from sewage using the PR4-positive *S. Typhimurium* LT2 as the host (Accesion PX306536). The phage showed 100% coverage and 97.5% nucleotide identity with the firstly reported Enterobacteria phage PRD1 in NCBI (NC_001421). This phage could lyse the PR4-positive *Salmonella, E. coli* and *P. aeruginosa*, whereas it could not lyse the RP4-negative bacterial counterparts. Phage pLT2 was isolated by using *S. Typhimurium* LT2 as host (Accession PX311067)and its genome showed 92% coverage and 96.1% nucleotide identity with *Salmonella* phage Felix O1 (NC_005282). pLT2 could lyse more than 40 different *Salmonella* serotypes in our lytic-spectrum testing (Supplementary table 1).

### Study design of chicken models

A total of 73 newly hatched Arbor Acres broilers (one day old) were included in this study. All broilers were acclimated to the new environment for 3 d. Then 65 broilers received an intragastric administration of 500μL of a PBS suspension containing 1.0 × 10^8^ CFU°mL^-1^ *gfp*-labelled RP4-positive *S. Typhimurium* LT2 every day for 7 days (day 4-day 10). The remaining 8 chicks were put in a separate sterile isolation chamber as uninfected controls. On day 11, five *Salmonella* infected chicks were randomly selected and euthanized to collect their intestinal content for metagenomic sequencing (pre-treatment control, Y). Then, the remaining 60 infected chicks were randomly divided into five groups (12 chickens per group) and placed in separate sterile isolation chambers. The groups were set as follows: LT2 phage treatment (group L; 5×10^9^ PFU/chick/day); PRD1 phage treatment (group P; 5×10^9^ PFU/chick/day); PRD1 and pLT2 phage cocktail treatment (group P+L; 1:1 phage mix, 5×10^9^ PFU/chick/day); Ampicillin treatment (group A; approximately 1 mg/chick/day); and an untreated control (group N). Phage treated chicks received the phage dissolved in PBS (500μL of 10^10^ PFU/mL phage stock)via intragastric administration continuously for 3 d (day 11 to day 13). For antibiotic treatment, the ampicillin (100 μg/mL) was administrated continuously via drinking for two weeks (day 11 to day 24).

On day 11,18, 25 and 35, three chickens from each treated group and untreated controls were euthanized to collect their intestinal content. RP4-positive *S. Typhimurium* LT2 were isolated from these intestinal contents on Bismuth Sulfite Agar supplemented with Ampicilin (100 μg/mL) plus chloramphenicol (50 μg/mL) and nalidixic acid (500 μg/mL). qPCR was used to determine the absolute abundance of RP4 and the administered phages in chicken intestinal contents at different time points (see supplementary methods for details of qPCR). We also collected gut contents for metagenomic sequencing from euthanized chickens that were uncolonized controls (day 11, n=5; day 35, n=3) and pre-treatment control chickens that were colonized with *Salmonella*, but not treated (day 11, n=5).

All broilers were provided with free access to feed and water throughout the 35-day trial. Both feed and water were subjected to PCR-based *Salmonella*-negative testing and sterilization treatment. The animal study received approval in adherence to the guidelines for the care and utilization of laboratory animals by the Jiangsu Academy of Agricultural Sciences (IACUC-AE-2023-01-020).

### Metagenomic sequencing and analysis

Metagenomic DNA was extracted from the chicken intestinal contents using DNeasy PowerSoil Pro Kit (47016; Qiagen, Germany) and sequenced using the MGI DNBSEQ-500 platform with a paired-end (PE) read length of 150 bp. Afterward, more than 10 Gb of raw reads underwent QC and host sequence filtering using Trimmomatic (v0.33) and BMTagger (v2.2.4) to eliminate adapter sequences, low-quality sequences, and contaminating chicken DNA. The clean reads were then assembled using MEGAHIT (v1.1.2), after which contigs of less than 500 bp were filtered. MetaBAT2 was used to calculate the coverage of all contigs and perform metagenomic binning to obtain metagenome-assembled genomes (MAGs) that were taxonomically equivalent to microbial strains. The taxonomic annotation for bins was inferred using GTDB-Tk (v. 2.2.0) with default parameters based on the Genome Taxonomy Database. Open reading frames (ORFs) of the assembled contigs were predicted using Prodigal (v. 2.6.3) in meta mode. Taxonomic classification of the intestinal microbiota was performed using Kraken2, and Bracken was used to calculate microbial diversity and abundance. In addition, reads were aligned to their respective assemblies using bowtie2 to quantify the abundance of contigs, ARGs, and MGEs. This was followed by normalization using the transcripts per kilobase million (TPM) metric, calculated by SAMtools.

To analyse changes in the chicken microbiome under different treatments, principal coordinates analysis (PCoA) based on Bray–Curtis distances was performed using the ape (v5.7-1) and vegan (v2.6-4) packages in R (v4.3.1). Spearman rank correlations were computed between the relative abundance of each family and the first two principal coordinates. The correlation matrices were visualized to highlight correlations between the relative abundance of each family and the first two principal coordinates. Trajectory analysis was then conducted to compare differences in size, direction, and shape of community change, using the first six principal coordinates, which together explained ∼60% of the variance in the relative abundance dataset. Statistical testing of trajectories was adapted from previously described approaches.

To identify RP4 variants, sequencing reads were mapped to the parental RP4 plasmid reference using BWA-MEM (v0.7.17). Duplicate reads were removed using Picard (v3.1.0). Coverage profiles were generated with BEDTools (v2.31.0). Variants were called from BAM files using SAMtools (v1.18) and VarScan (v2.4.6), with thresholds of ≥5% allele frequency, ≥5× coverage at the SNP site, and *P* < 0.05. Variants were subsequently annotated using SnpEff (v5.2).

### Data availability

Raw data of the metagenome sequences generated in this study are accessible through the NCBI Sequence Read Archive under the BioProject accession number BioProject ID PRJNA1327878.

## Results

### PRD1 treatment supresses *Salmonella* and the RP4 plasmid in chicken gut microbiomes

To assess the impact of phage treatments, we measured the abundance of the *Salmonella* donor strain and the RP4 plasmid in phage treated chickens relative to untreated controls (Figure 2). Phage treatments that contained PRD1 strongly reduced abundance of the *Salmonella* donor strain relative to untreated controls, driving its clearance from the gut microbiome by day 35 (Figure 2A). Treatment with pLT2 alone initially suppressed the abundance of the *Salmonella* donor strain relative to untreated controls, but this effect was transient and failed to clear *Salmonella* from the gut microbiome. In contrast, treatment with ampicillin was associated with the stable maintenance of *Salmonella* in the gut microbiome.

**Figure 2:**
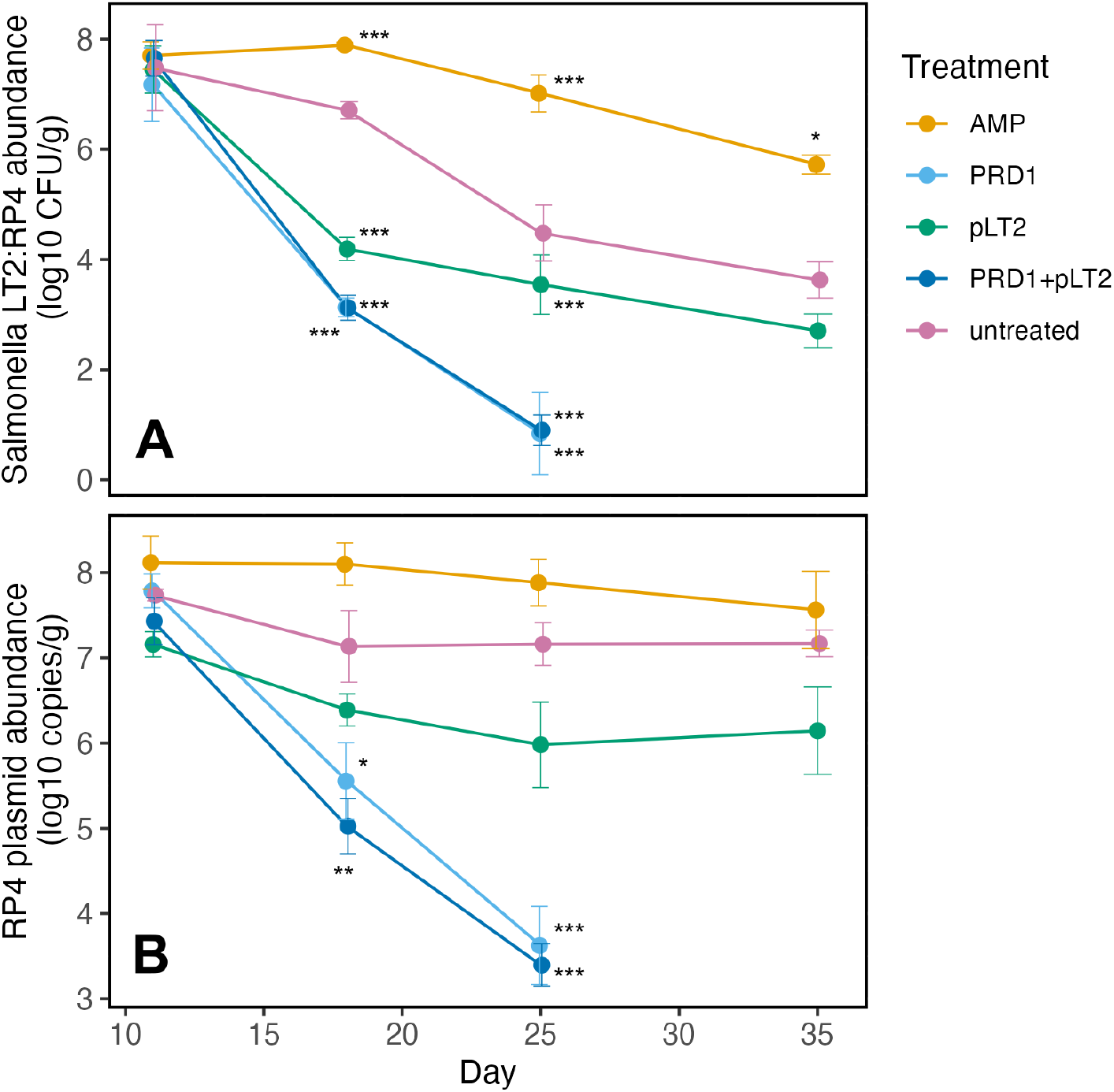
PRD1 treatments eliminates AMR from the chicken gut microbiome. Chickens colonized with a host strain of *Salmonella* carrying the RP4 plasmid were treated with PRD1 and pLT2, both alone and in combination, or with ampicillin, which selects for the RP4 plasmid. Chickens were sacrificed (n=3/treatment/time point) at regular intervals until day 35. (A) The density of the *Salmonella*:RP4 host strain was determined by selective plating, with a lower limit of detection of 10 CFU/g. (B) The density of the RP4 plasmid in the gut microbiome was determined by qPCR, with a lower limit of detection of 10^3^ plasmid copies/g. Plotted points show means (+/-s.e.m;n=3) and densities were compared to untreated control chickens (n=3/time point) at each day using Dunnett’s test (* P<.05;** P<.01; *** P<.001).

The RP4 plasmid was maintained in untreated controls at comparable levels to ampicillin treated chickens, highlighting the ability of RP4 to persist in the absence of antibiotic treatment (Figure 2B). RP4 transferred into the background microbiota at a high rate in untreated and pLT2 treated chickens, evident from log-fold higher RP4 than *Salmonella* abundance from day 25 onwards. In contrast, treatments containing PRD1 strongly reduced RP4 abundance relative both to untreated controls and pLT2 treatment. The magnitude of this effect was impressive: the difference in RP4 abundance between PRD1 treated chickens and untreated controls was at least 1,000x by day 25, and by day 35 RP4 was eliminated by PRD1 from the chicken gut microbiome. These quantitative measures of RP4 dynamics were supported by directly visualizing GFP production in samples of chicken intestinal contents by confocal fluorescent microscopy (Figure S1).

Together, our data show that PDP therapy was highly effective, driving the complete elimination of both *Salmonella* and the RP4 plasmid from the chicken microbiome within 35 days. This clearance occurred through a combination of PRD1 directly killing the *Salmonella* donor strain and blocking RP4 transfer into the background microbiota.

The densities of both phage declined over time, suggesting that phage successfully depleted susceptible host cells (Figure 3). Using phage cocktails generates the potential for competition between phage to infect bacterial host cells^37,38^. However, the densities of each phage did not differ between the phage cocktail and the corresponding single phage treatment, suggesting that an appreciable fraction of PRD1 infections were of transconjugant cells carrying the RP4 plasmid in the combined phage treatment.

**Figure 3:**
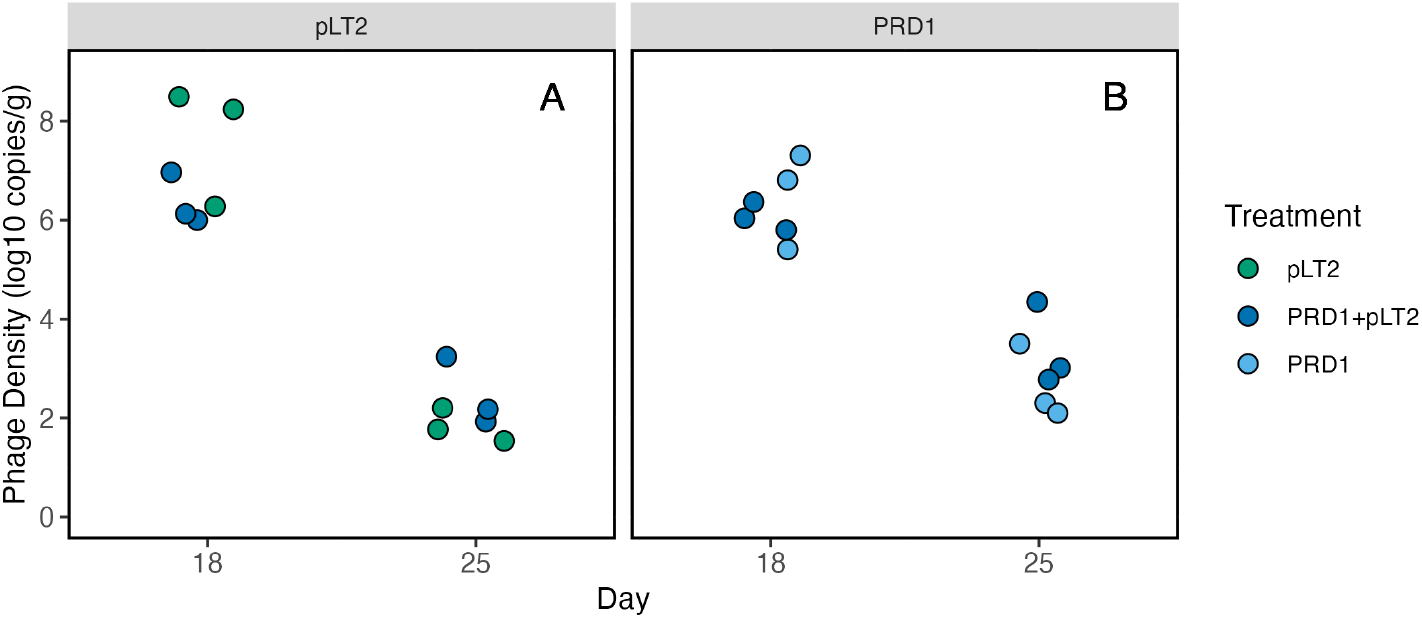
Phage population dynamics. Plotted points show the density of phage pLT2 (A) and PRD1 (B) in the chicken gut intestine, as assessed by qPCR. Each data point represents a single chicken. Phage densities at day 11 and day 35 were below the detection limit of 10 copies/g. Phage densities were not decreased by the presence of competing phage, as assessed by ANOVAs including a main egects of treatment (ie monotherapy or phage cocktail) and day (*P*>.5).

### PRD1 treatment has minimal impact on the composition of the chicken gut microbiome

Given the narrow host range of phage, phage therapy is expected to have minimal collateral impacts upon gut microbiota composition^39^. However, the suppression of the target pathogen by phage can result in indirect effects upon untargeted taxa that interact ecologically with the pathogen^40^. To test for broader impacts of phage treatment on the microbiota, we used metagenomics to compare the microbiome composition of phage treated chickens with both untreated controls and chickens that were treated with ampicillin as a positive control for microbiome perturbation (Figure 4A, B, C and D).As expected, treatment with ampicillin was associated with a significant perturbation to the microbiome relative to untreated controls, in terms of more change in microbiome composition over time and greater divergence of microbiome composition at day 35 (Figure 4C and D, permutational MANOVA, 10,000 permutation, treatment x time interaction, *F* = 0.058, *p* = 0.014, pairwise absolute differences in path distance, A-N: *Z* = 0.591, *p* = 0.007). In contrast, phage treatments were minimally disruptive to microbiome composition, showing similar dynamics of microbiome composition as untreated controls over time (pairwise absolute differences in path distance, P-N: *Z* = 0.038, *p* = 0.856; PL-N: *Z* = 0.207, *p* = 0.341; L-N: *Z* = 0.105, *p* = 0.626). Chicken gut microbiomes treated with PRD1 or both phages were indistinguishable from uninfected controls by day 35, whereas treatment with either ampicillin or pLT2 alone showed sustained microbiome disruption (Figure 4C).

**Figure 4.**
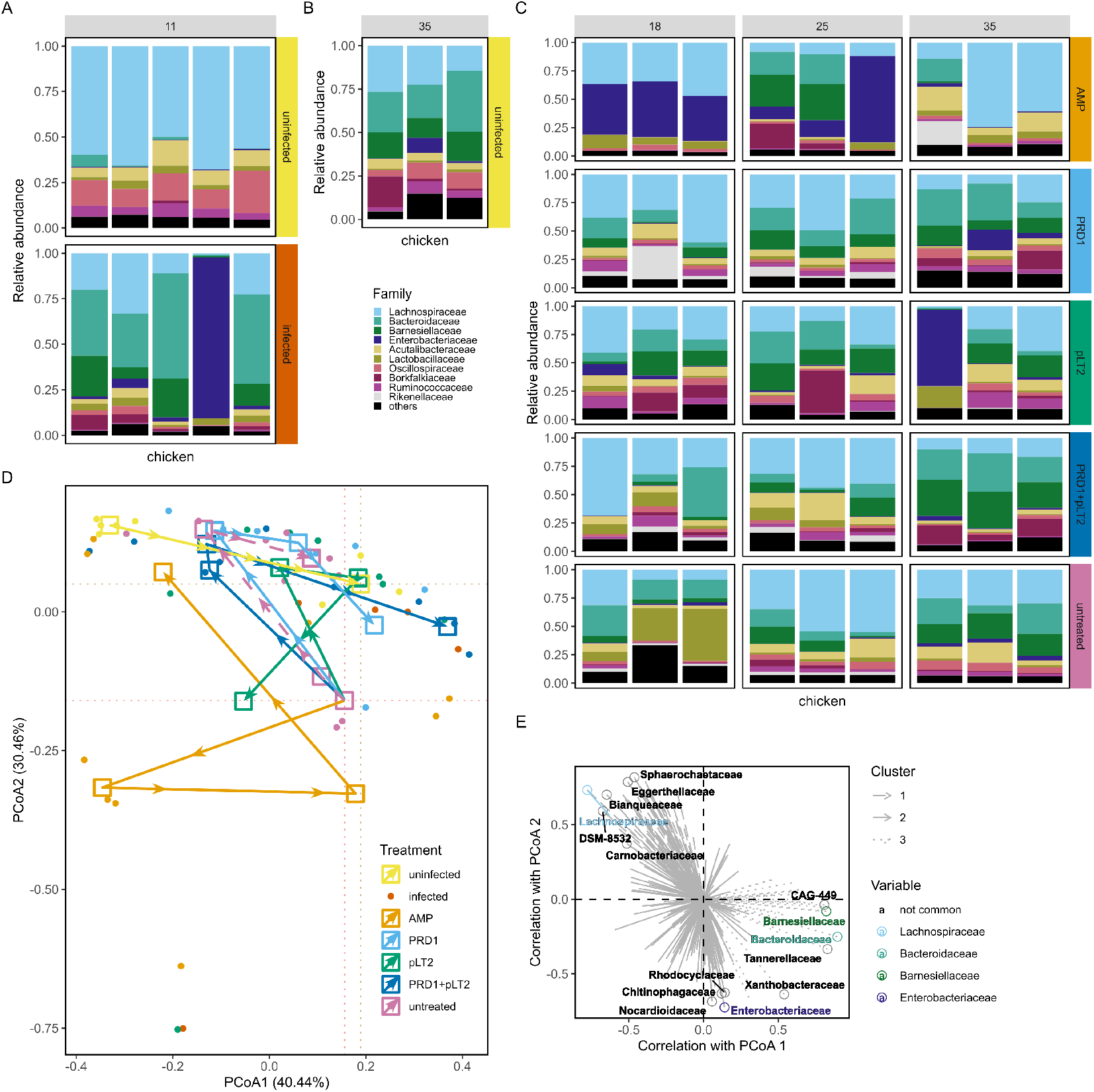
Changes in microbial composition in chicken gut microbiome. The 10 most abundant bacterial families in chicken gut microbiomes were observed at multiple time points: (A) pre-(upper panel) and post-*Salmonella* infection (lower panel) on day 11; (B) in uninfected chickens at day 35; and (C) in infected chickens across days 18, 25, and 35. (D) Treatment-induced microbiome shifts were quantified by comparing mean compositional changes relative to uninfected controls characterized by Principal Coordinate Analysis (PCoA) using Bray-Curtis distance matrix of relative abundance at the family level. Microbial compositions of individual chickens are shown as single dots. (E) Family-level abundances were correlated with the first two principal coordinates (PCs) of PCoA and clustered into three distinct groups, with the five families showing the highest magnitudes in each cluster being labelled.

**Figure 5:**
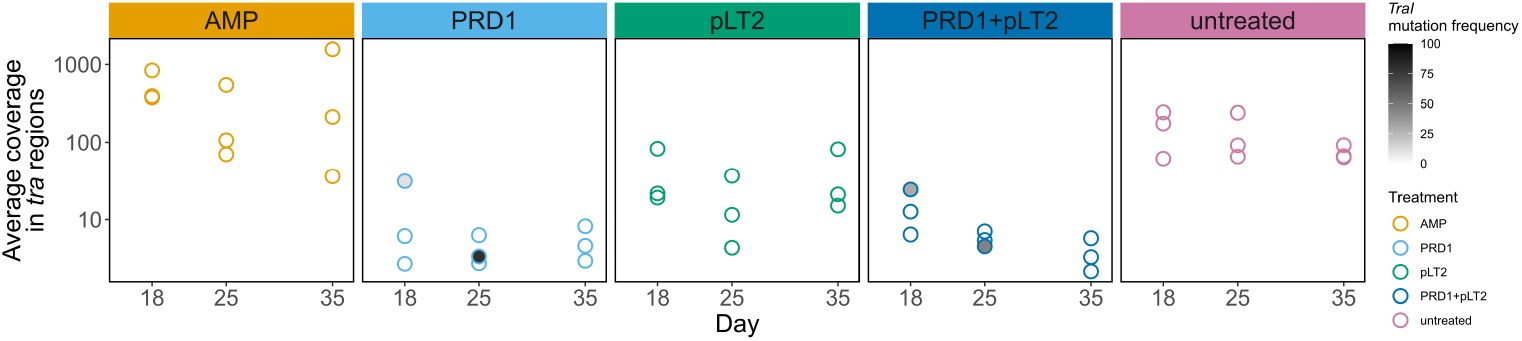
Selection for *tra* variants. Plotted points show the average depth of coverage of the RP4 *tra* region from metagenomic sequencing of *Salmonella* colonized chickens. Shading indicates the frequency of the *traI* Arg524Gly polymorphism, which was detected in a significant minority of PRD1 treated chickens (n=4/18) compared to chickens that were not treated with PRD1 (n=27) as judged by Fisher’s exact test (Odds ratio= ∞, *P* =.0205). Note that this is a conservative test because the low coverage of PRD1-treated chickens was associated with a reduced likelihood of detecting polymorphisms.

### PRD1 selects for mutations in the RP4 conjugative region

RP4 adapts to PDR1 in the lab through mutations affecting the plasmid’s conjugative apparatus^15,41^. We compared plasmid evolution between treatments by mapping metagenomic sequencing reads to the RP4 plasmid genome. To reduce noise arising from mismapping of other genetic elements endogenous within the chicken gut microbiome, we focused on calling variants within the conjugation regions of RP4, which we expected to be less represented in the chicken gut microbiome compared to RP4 cargo genes (ie IS elements, antibiotic resistance genes).

Using this approach we detected very little evidence for *in vivo* evolution of the RP4 conjugation region. A missense polymorphism (Arg524Gly) in *TraI* was detected in 4/18 chickens that were treated with PRD1. This polymorphism was not detected in the 27 colonized chickens that were not treated with PRD1, suggesting this mutation is a parallel evolutionary response to PRD1 treatment. TraI is an essential component of the relaxosome complex^42^, and the Arg524Gly polymorphism occurs in a conserved helicase domain of the protein^43,44^, suggesting that this mutation may inhibit conjugation. However, the presence of the TraI polymorphism did not appear to negatively impact PRD1 efficacy, as it was not associated with an increased prevalence of either *Salmonella* or the RP4 plasmid (P>0.05).The only other protein-altering mutation in the conjugative region was found in a single PRD1 treated chicken and an untreated chicken (VirB4 Val430Leu; n=2), suggesting that this mutation may be an adaptation to the chicken gut and not a response to PRD1 *per se*.

## Conclusion

The key finding of this study is that treatment with the PRD1 pilus dependent phage efficiently suppresses *Salmonella* and prevents the dissemination of the RP4 plasmid to the gut microbiome. As such, PRD1 treatment offers a clear advantage over conventional phage therapy targeting only *Salmonella*. This is because only killing the target pathogen still allows the RP4 plasmid to persist by transferring to the background gut microbiota, permitting maintenance and spread of the encoded AMR genes within the microbiome. These findings suggest that PDPs may be a particularly important tool for combatting bacteria carrying highly conjugative AMR plasmids that are rapidly transferred between commensal and pathogenic bacteria in the gut microbiome^8^. PRD1 can infect any bacteria carrying the RP4 multidrug resistance plasmid^22^, and this broad host range suggests that PDP therapy is likely to be especially useful in microbial communities that are hot spots of plasmid transfer, such as the gut microbiome. Importantly, however, PRD1 had no sustained off-target effects on the composition of the gut microbiome, suggesting that PDPs are unlikely to disrupt the microbiota at a community-wide scale.

Mutations linked to phage resistance are commonly observed following phage therapy, and can compromise treatment efficacy ^1,40,45^. The RP4 plasmid was present at a very high abundance at the outset of phage treatment, implying that the plasmid had a high underlying potential for adapting to the strong selective pressure imposed PRD1 treatment. However, metagenomic sequencing suggests that mutations in the conjugation region likely to cause resistance to PRD1 evolved rarely, and were not associated with treatment failure. How can we explain this apparent paradox? The RP4 plasmid successfully persisted in untreated chickens by efficiently transferring from the *Salmonella* donor strain to the gut microbiome (Figure 2). This result suggests that PRD1 treatment was successful because conjugation is essential for the RP4 plasmid to persist in the gut microbiome, limiting the potential for PRD1 resistance via mutations impairing conjugation to evolve. The essentiality of the conjugative apparatus in the gut environment may help to explain the convergent evolution of pilus targeting in diverse phage groups^17,21,22^, and the high rate of extinction of plasmids following the loss of conjugative genes^46^. A key challenge for future research will be to understand whether other AMR-associated plasmids depend on conjugation to persist in bacterial communities^8,16,47-51^ and the extent to which AMR can be suppressed in human and animal microbiomes by targeting conjugation^51-53^.

## Acknowledgements

This project received funding from UKRI (MR/W031361/1) and ICARS (200001), under the umbrella of the JPIAMR -Joint Programming Initiative on Antimicrobial Resistance. This work was supported by UKRI Frontiers Grant (EP/Y031067/1) to RCM and a BBSRC Engineering Biology Mission Award (BB/Y007743/1) and a BBSRC sLoLa grant (BB/X003051/1) to MAB.

## CRediT taxonomy

Jun Li: Experiment conducting and Formal Analysis

Taoran Fu: Formal Analysis, Visualization, Writing – review & editing

Xing ji: Experiment conducting and Formal Analysis

Ran Wang: Conceptualization and Formal Analysis

R.Craig MacLean: Conceptualization, Funding Acquisition, Formal Analysis, Visualization, Writing – original draft, Writing – review & editing

Michael A. Brockhurst: Conceptualization, Funding Acquisition, Writing – original draft, Writing – review & editing

Tao He: Conceptualization, Funding Acquisition, Writing – original draft, Writing – review & editing

