## Supplementary methods and figures for "Plasmid dependent phage eliminate pathogenic bacteria and antibiotic resistance plasmids from the chicken gut microbiome"

### Supplementary Materials

#### Supplementary Methods

##### 1. Protocols for construction of *gfp*-labelled RP4 plasmid

The *gfp* gene with promoter was cloned into RP4 plasmid by the homologous recombination method.

**1.1 Primer design**

Restriction site snaBI and SbfI in RP4 plasmid were selected for primer design. The primer sequences were shown in Table 1.

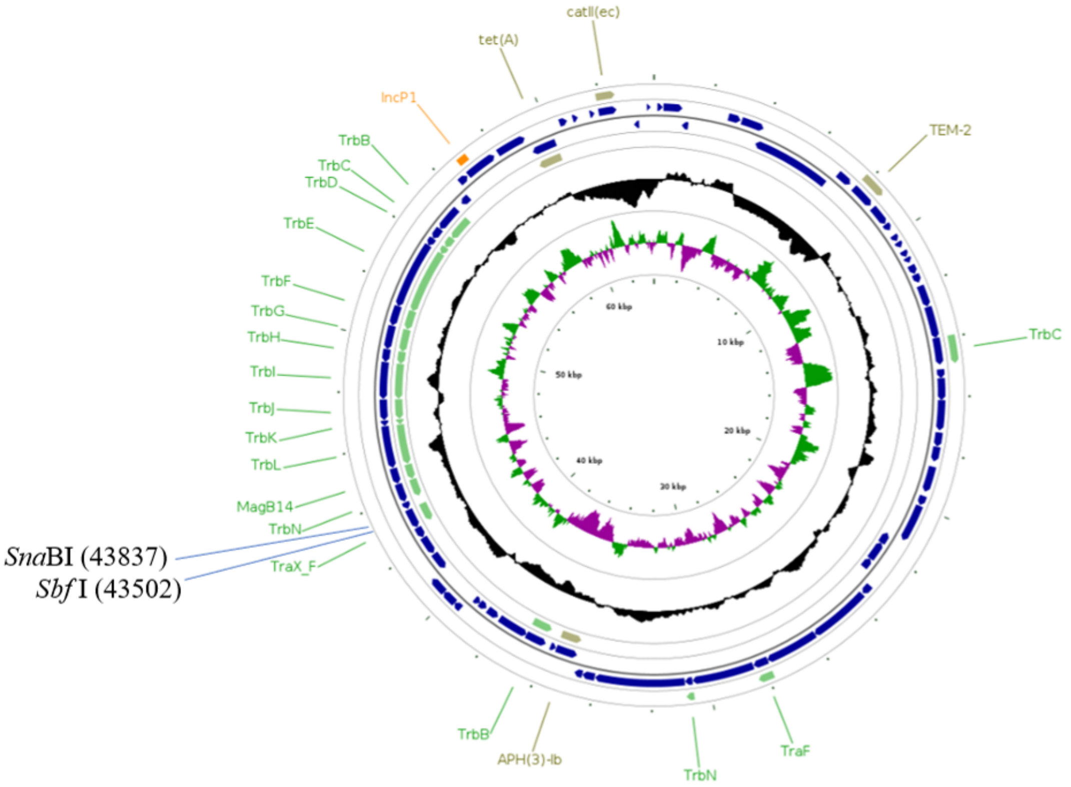


Fig. 1. RP4 plasmid map

Table 1. Primer Sequences

| Primers | Primer sequence (5'-3') |
|---|---|
| F1 | CCCCTTGGAGTAAGAACGC（**RP4 homologous sequence**）**CCTGCAGG**（SbfI restriction site）ATAAAAAACGCCCGGCGGCAACC (gfp terminator homologous sequence) |
| R1 | CAGCCTCCGCAGCTCGGGT（**RP4 homologous sequence**）**TACGTA**（snaBI restriction site）GGCGCGCCCCTCCTTGAC（gfp promotor homologous sequence) |

**1.2** The *gfp* gene with the promoter and terminator was synthesized by one Bio-technology company and the sequences are as followings:

(Promotor)GGCGCGCCCCTCCTTGACACTGAATTTAGCATGTGATATAATTAACTTAAT
ATTCTACCCAAGCTTATAAAAGAGCACTGTTGGGCGTGAGTGGAGGCGCCGGAAA
AAAGCATCGAAAAAAGGAGGAAAAAAA**ATGGTATCAAAAGGGGAAGAGCTGT
TCACAGGCGTTGTACCAATTTTGGTGGAGCTGGACGGTGATGTTAATGGCCA
CAAGTTTTCCGTGAGCGGCGAGGGTGAAGGTGACGCAACGTATGGCAAACTT
ACGTTAAAGTTCATTTGCACCACGGGCAAGTTGCCAGTTCCTTGGCCGACAT
TAGTGACCACTCTTACATACGGAGTTCAATGTTTCTCACGTTACCCGGATCAC
ATGAAACAGCACGATTTCTTCAAGTCAGCTATGCCCGAGGGTTATGTTCAGGA
GCGCACGATCTTCTTCAAAGACGATGGTAATTACAAAACACGCGCGGAAGTG
AAGTTCGAGGGCGATACACTTGTTAATCGTATTGAATTAAAAGGGATCGACTT
TAAAGAAGACGGGAATATCTTGGGTCACAAATTAGAATATAATTATAACAGTCA
CAACGTATACATTATGGCGGACAAACAGAAAAACGGAATCAAAGTGAACTTTA
AAATTCGTCACAACATTGAGGATGGGTCCGTACAGCTTGCAGATCACTATCAA
CAAAACACTCCTATCGGCGACGGACCTGTTCTGTTACCGGACAACCACTATCT
TAGTACACAGTCAGCATTAAGCAAAGACCCGAACGAGAAGCGTGACCACATG
GTGTTGCTTGAGTTCGTGACTGCGGCAGGCATTACACTGGGTATGGATGAAC
TGTACAAGTAA**TCAGTAGATCTCTGCAGTCGCGATGATTAATTAATTCAGAACGCT
CGGTTGCCGCCGGGCGTTTTTTAT(Terminator)

**1.3** The *gfp* gene with the promoter and terminator was amplified by PCR and the PCR reaction system and reaction procedures are shown in Table 2 and Table 3. The PCR amplicons were purified and recovered by Gel DNA Purification Kit.

Table 2. PCR reaction system for *gfp* gene

| Composition | Volume (μL) |
|---|---|
| 2 ×PCR Mix (Dye Plus) | 10 |
| Primer F | 1 |
| Primer R | 1 |
| $ddH_2O$ | 6 |
| *gfp* gene template | 2 |

Table 3. PCR reaction procedures for *gfp* gene

| Reaction temperature | Time | Number of reaction |
|---|---|---|
| 95°C | 5 min | 1 |
| 95°C | 15 s | |
| 60°C | 30 s | 32cycles |
| 72°C | 1 min | |

| | | |
|---|---|---|
| 72°C | 10 min | 1 |

**1.4** The RP4 plasmid was digested by snaBI and SbfI at 37°C overnight. The enzymatic digestion reaction system are shown in Table 4. The digestion product was purified and recovered by DNA Purification Kit.

Table 4. Enzymatic digestion reaction system

| Composition | Volume (μL) |
|---|---|
| SnaBI | 1 |
| SbfI | 1 |
| 10×buffer | 5 |
| RP4 plasmid DNA | 1 |
| $ddH_2O$ | Up to 10μL |

**1.5** Homologous recombination between linearized RP4 plasmid and the GFP amplicon at 50°C for 15 min using Seamless Cloning Kit. The reaction system are shown in Table 5.

Table 5. Seamless cloning reaction system

| Composition | Volume (μL) |
|---|---|
| linearized RP4 plasmid | 3 |
| GFP amplicon | 2 |
| 2×Seamless Cloning Mix | 10 |
| $ddH_2O$ | up to 20 |

**1.6** The recombined RP4 plasmid was transformed into the *E. coli* DH5a and *S. typhimurium* LT2 competent cells by electroporation. The transformants were selected by Ampicillin and confirmed by PCR detection of the *gfp* gene. The *gfp*-labelled RP4-positive *S. typhimurium* LT2 was further confirmed by observation under UV and fluorescence confocal microscopy. (Fig. 2)

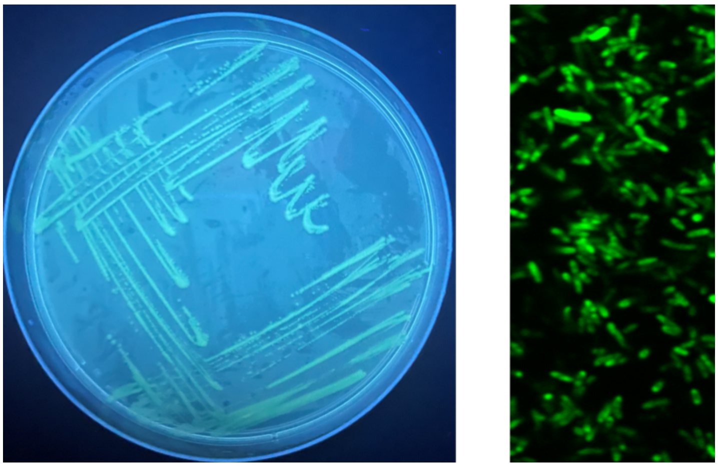

Fig. 2. The *gfp*-labelled RP4-positive *S. typhimurium* LT2 under UV and fluorescence confocal microscopy

#### 2. Protocols for qPCR detection of RP4 plasmid, PRD1 and LT2 phages

**2.1** The qPCR primers are designed on the conservative region of the targets and the primers are listed in Table 6.

Table 6. Primer Sequences

| Target | Primers | Primer sequence (5'-3') |
|---|---|---|
| RP4 | gfp-F | CGACGGACCTGTTCTGTTAC |
| plasmid | gfp-R | TCAAGCAACACCATGTGGTC |
| PRD1 | PRD1-F | TACTCCGGCCTTGAGTAAGTAG |
| phage | PRD1-R | TTGCTTTGGGATATTCACGCG |
| pLT2 | pLT2-F | ACGCATACTGTTGGTGGTCA |
| phage | pLT2-R | TGTTGGTGATTGCCTGTGGT |

**2.2** The standard curves were determined for RP4 plasmid ( log[template]=-3.611*CT+40.45), PRD1 phage (log[template]=-3.229*CT+35.13) and pLT2 phage (log[template]=-3.173*CT+36.52), respectively. The linear range of standard curves (R>0.99) were $10^3$~$10^9$ copies/g for RP4 plasmid, $10^1$~$10^{10}$ PFU/g for PRD1 phage and $10^1$~$10^9$ PFU/g for the pLT2 phage, respectively.

**2.3** qPCR were performed on Roche LightCycler 96. For RP4 plasmid detection, the whole DNA of chicken intestinal content were extracted using DNeasy PowerSoil Pro Kit. For phage detection, the intestinal contents were mixed thoroughly with 10-fold volume of PBS and centrifuged to collect the supernatant, and the supernatant was used as DNA template for qPCR. The reaction systems and reaction conditions were shown in Table 7 and Table 8.

Table 7. qPCR reaction system

| Composition | Volume (μL) |
|---|---|
| 2 × Taq Pro Universal SYBR Master Mix | 10 |
| Primer F | 0.4 |
| Primer R | 0.4 |
| ddH2O | 8.2 |
| DNA or phage supernate | 1 |

Table 8. qPCR reaction program

| Reaction temperature | Time | Number of reaction |
|---|---|---|
| 95°C | 30 s | 1 |
| 95°C | 10 s | 40 cycles |
| 60°C | 30 s | |
| 95°C | 15 s | 1 |
| 60°C | 1 min | 1 |
| 60°C | 15 s | 1 |

**2.4** The Ct values of the samples were substituted into the standard curve formula and calculated for the absolute abundance of the targets.

#### Supplementary figure

**Figure S1. Direct visualization of plasmid abundance.**

Samples of intestinal contents from the n=3 chickens from each treatment/time point combination were combined together as a single sample. GFP production from the RP4 plasmids was visualized using a fluorescence confocal microscope, and the scale bar shows 29 μm for each panel.

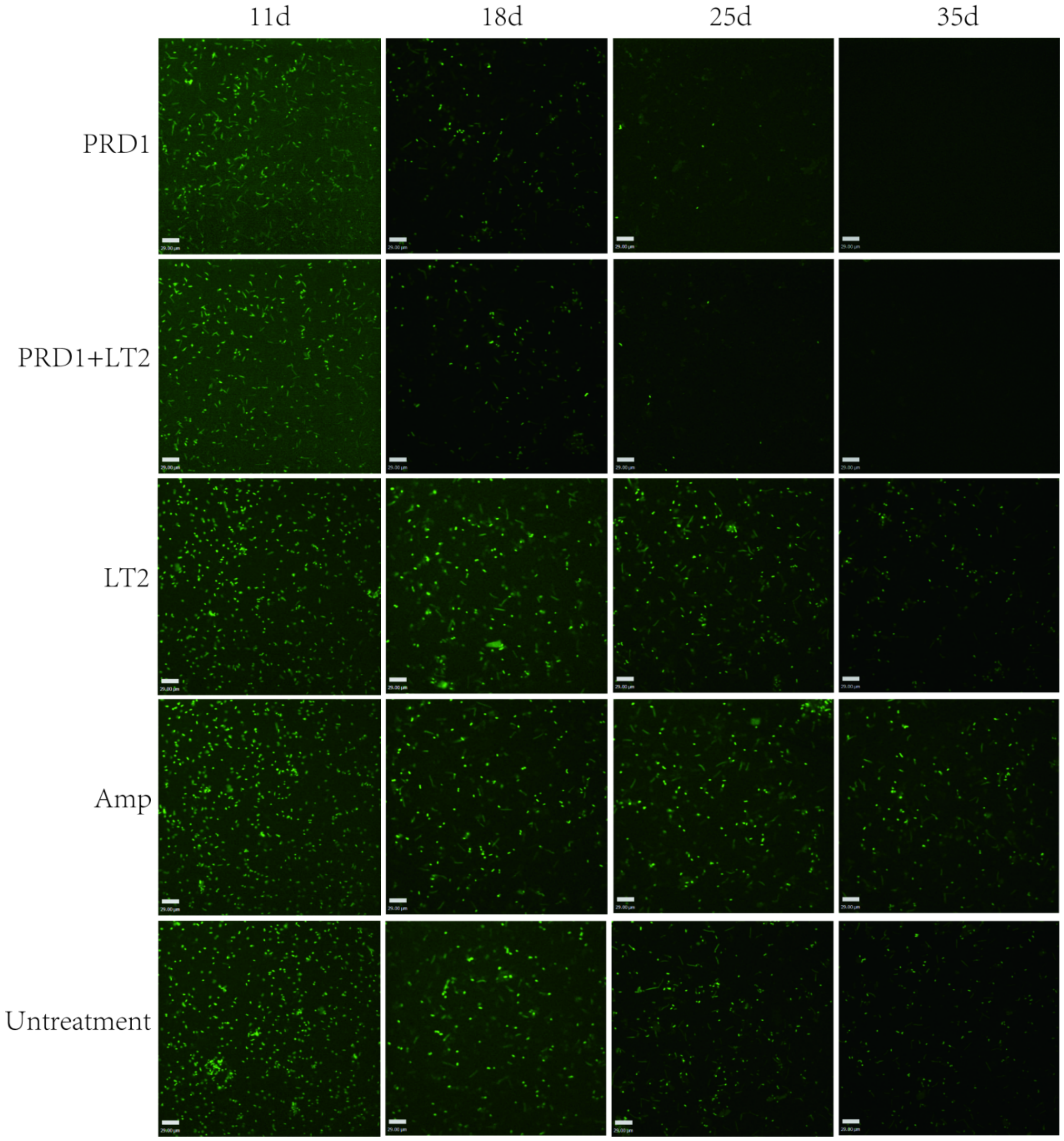

#### Supplementary table

**Table S1.** The lytic spectrum of phage pLT2 used in this study.

| **Strain designation** | **Serotype** | **Lytic ability** |
|---|---|---|
| p159-1 | Aberdeen | + |
| XC-5-1Z | Agona | + |
| ws010 | Agoueve | + |
| L35 | Albany | + |
| 11 | Bareilly | + |
| XC-5-JC | Braenderup | + |
| 19SC14RX37 | Corvallis | + |
| FS-1-10Z | Derby | + |
| 186363 | Dublin | + |
| L3 | Enteritidis | + |
| FS-1-7Y | Fanti | + |
| 23S20 | Give | + |
| FT-6-5JT | Hadar | + |
| z91 | Huddinge Lerum | + |
| L80 | Indiana | + |
| 23S50 | Infantis | + |
| 23S48 | Javiana | + |
| HD-3-JX | Kedougou | + |
| SJS-2-JX | Kentucky | + |
| SM22KDE | Kottbus | + |
| Li-S1 | Litchfield | + |
| SY-1-2IN | London | + |
| FT-3-JC | Mbandaka | + |
| z90 | Meleagridis | + |
| CP-1-3JC | Muenster | + |
| 23S37 | Newport | + |
| 23S12 | Pomona | + |
| PG-1-20JX | Portanigra | + |
| 192093 | Potsdam | + |
| P001-4-2 | Pullorum | + |
| CY-1-6Z | Rissen | + |
| SJS-1-JC | Saintpaul | + |
| 366364 | salamae | + |
| DX-4-JC | Schwarzengrund | + |
| Sen-1 | Senftenberg | + |
| stan-1 | Stanleyville | + |
| SM31KDE | Tennessee | + |

| | | |
|---|---|---|
| tho-1 | Thompson | + |
| **LT2** | Typhimurium | + |
| 23S6 | Uganda | + |
| z35 | Weltevreden | + |

Note: + indicates bacterial lysis by using the agar spotting test.
